# Neural and behavioral signatures of policy compression in cognitive control

**DOI:** 10.1101/2025.05.06.652533

**Authors:** Shuze Liu, Atsushi Kikumoto, David Badre, Samuel J. Gershman

## Abstract

Making context-dependent decisions incurs cognitive costs. Cognitive control studies have investigated the nature of such costs from both computational and neural perspectives. In this paper, we offer an information-theoretic account of the costs associated with context-dependent decisions. According to this account, the brain’s limited capacity to store context-dependent policies necessitates “compression” of policies into internal representations with an upper bound on code length, quantified by an information-theoretic measure (policy complexity). These representations are decoded into actions by sequentially inspecting each bit, such that longer codes take more time to decode. When a response deadline is imposed, the account predicts that policy complexity should increase with the deadline. Higher policy complexity is associated with several behavioral signatures: (i) higher accuracy; (ii) lower variability; and (iii) lower perseveration. Analyzing data from a rule-based action selection task, we found evidence supporting all of these predictions. We further hypothesized that complex policies require higher neural dimensionality (which constrains the code space). Consistent with this hypothesis, we found that policy complexity correlates with a measure of neural dimensionality in a rule-based decision task. This finding brings us a step closer to understanding the neural implementation of policy compression and its implications for cognitive control.

## Neural and behavioral signatures of policy compression in cognitive control

A prominent hallmark of human cognition is our ability to select actions appropriate for the current context. The study of cognitive control seeks to understand this core capability (Badre, 2024; Fan, 2014; Shenhav et al., 2017). A growing body of research has highlighted the cognitive costs incurred by such control processes. These costs are evident in human tendencies to avoid tasks that demand greater cognitive control (Kool et al., 2010; Sayalı et al., 2023; Shenhav et al., 2017), and increase control under greater reward incentives (Krebs and Woldorff, 2017; Umemoto and Holroyd, 2015; Shenhav et al., 2017).

The ubiquity of cognitive costs has prompted researchers to probe their underlying mechanisms. Computational models have proposed a range of explanations, from metaphorical force fields to theories of limited mental resources and reward-based accounts of effort allocation (Botvinick and Braver, 2015; Kruglanski et al., 2012; Shenhav et al., 2017). In parallel to the computational perspectives above, neuroimaging studies have localized control-related signals in the brain, including but not limited to the anterior cingulate cortex (ACC) and lateral prefrontal cortex (LPFC) (Badre, 2008; Koechlin and Summerfield, 2007; McGuire and Botvinick, 2010). While these perspectives are in principle reconcilable (Botvinick and Cohen, 2014), their integration has been limited by the absence of a unifying computational resource formulation—one that can both reconcile these diverse theories and be empirically tested.

In this paper, we take a step toward such a unifying perspective by refining an information-theoretic framework of cognitive control. Building on previous theoretical and empirical work, we explore how this perspective can bridge behavioral and neural accounts of cognitive effort. The idea of connecting information theory to cognitive control is not new: early cognitive control studies on the PFC proposed a hierarchical architecture along a rostro-caudal axis, with anterior regions supporting higher-order, more abstract-level control. These ideas were formalized using information-theoretic measures including mutual information and conditional entropy, interpreted as measures of remaining uncertainty within the hierarchy (Koechlin and Summerfield, 2007; Badre, 2008). While these information-theoretic measures have the benefit of being domain-general and thus generalizable across tasks, the above line of work did not specify how such information-theoretic constructs may reconcile the various cognitive cost formulations at the computational level, or be tested using behavioral data.

More recently, the importation of rate-distortion theory (Cover, 1999) into cognitive science has filled this gap. By leveraging its formalism of constrained optimization—balancing reward against channel information rate (Sims, 2016; Tishby and Polani, 2010; Zenon et al., 2019; Lai and Gershman, 2021)—this line of work has offered normative models of human decision-making that naturally connect the resource and reward-based views of cognitive control (Lai and Gershman, 2021). Additionally, the identified solution bears similarities with Kullback-Leibler (KL) regularized control and the way it penalizes deviations from a default policy (Todorov, 2009), enabling closer connections to the force field perspective. The resulting *policy compression* framework has been empirically supported in contextual bandit tasks (Gershman, 2020; Lai and Gershman, 2024; Liu et al., 2024; Liu and Gershman, 2025), and has begun to find traction in neuroscience (Gershman and Lak, 2025). While some connections to cognitive control—such as task switching and the overriding of habits—have been established (Zenon et al., 2019), few empirical studies have quantitatively assessed the framework’s predictions for cognitive control.

In this paper, we introduce the policy compression framework as a normative theory of context-dependent decision making, articulating its implications for three major issues in cognitive control: the formulation of cognitive cost as the mutual information between context and action (policy complexity), the emergence of perseverative behavior toward default actions, and the link between control and response times. We also propose an algorithmic-level implementation of the framework based on entropy coding, connecting the framework to recent findings on neural representational dimensionality and their role in cognitive control (Rigotti et al., 2013; Bernardi et al., 2020; Kikumoto et al., 2024a). In particular, the framework predicts that representational dimensionality should scale with policy complexity, under the assumption that more complex policies consume more representational resources. To evaluate these predictions, we analyze the behavior and neural activity of human participants performing a rule-based action selection task under varying response deadlines.

## The policy compression framework

Here we describe the proposed policy compression framework in detail. We first outline its foundation in rate-distortion theory, which formalizes context-dependent action selection as a reward optimization problem under information-theoretic constraints. We then elaborate on the framework’s implications for cognitive control, highlighting how it accounts for perseverative behavior during action selection and elucidates the links between state-dependent policies, response times, and representational dimensionality.

The nervous system operates under numerous constraints (Shenhav et al., 2017). These constraints have been formalized at multiple levels of analysis, ranging from computational-level accounts of sampling costs and computational complexity (Bossaerts et al., 2019; Zhou et al., 2024; Bossaerts et al., 2019; Vul et al., 2014; Ma et al., 2014), to physiological-level costs of interference and neural metabolism (Musslick et al., 2016; Gailliot and Baumeister, 2007). Here, we specifically focus on the influence of channel capacity, the maximum information that can be transmitted across a noisy channel (Shannon, 1948), on decision-making processes.

The framework we propose is an application of rate-distortion theory to action selection. Rate-distortion theory prescribes how to construct an optimal channel that minimizes some notion of error (the distortion), or maximizes reward, subject to a constraint on the information transmission rate (i.e., an information bottleneck) (Cover, 1999). The utility of rate-distortion theory lies in its task-general formulation of cognitive constraints. Beyond action selection (Lai and Gershman, 2021, 2024; Liu et al., 2024; Liu and Gershman, 2025), it has been successfully applied to various cognitive processes including visual working memory (Sims et al., 2012; Sims, 2015; Jakob and Gershman, 2023), perception (Gershman and Burke, 2023), intertemporal decision-making (Gershman and Bhui, 2020), and cognitive abstraction formation (Genewein et al., 2015). Previous works have linked information theory to various facets of cognitive control, from task-switching costs (Zenon et al., 2019) to the hierarchical organization of executive function (Koechlin and Summerfield, 2007; Badre, 2008). These studies support a *modulatory* view of control, in which top-down processing of higher-level contextual cues enhances or inhibits lower-level stimulus-response associations to guide behavior (Badre et al., 2021; Aron, 2007; Goghari and MacDonald III, 2009). Such connections enable us to apply rate-distortion theory to cognitive control through the lens of policy compression, which we will elaborate on below.

We assume a contextual decision-making setup, in which the agent selects actions based on some environmental state to maximize reward (Koechlin and Summerfield, 2007; Badre, 2008; Zenon et al., 2019; Kikumoto et al., 2022, 2024a). In this paper we will use “context” and “state” interchangeably. The agent’s policy π(*a*|*s*) is a probabilistic mapping from states *s* ∈ S to actions *a* ∈ A. Here we make the simplifying assumption that all contextual information is encapsulated into a (potentially high-dimensional) state *s* that informs action selection (Figure 1A).

**Figure 1.**
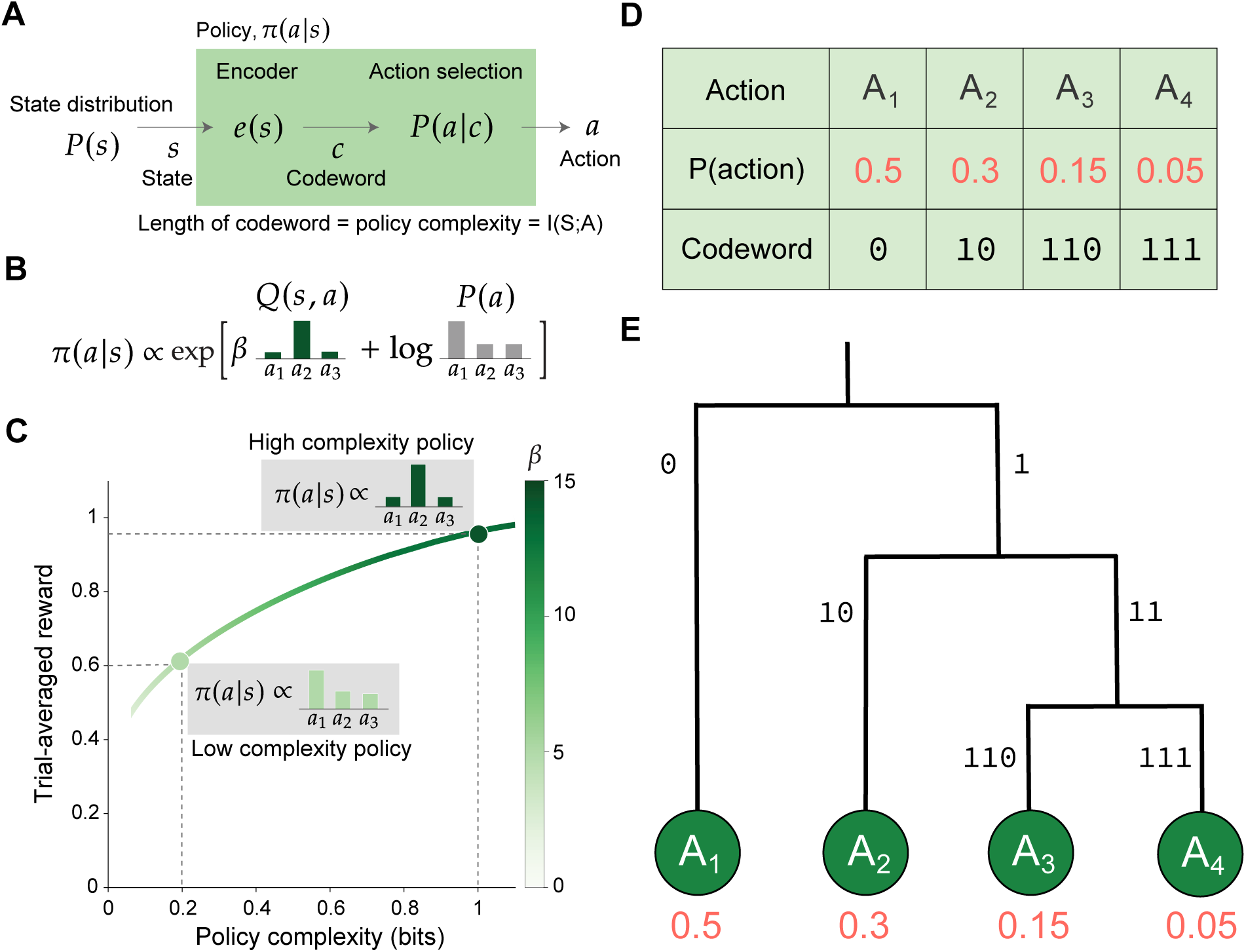
The policy compression framework and optimal entropy codes. **(A)** Context-dependent action selection as an information channel. The context/state is encoded as a codeword and subsequently decoded to reveal its policy-assigned action. The information channel is capacity-limited, which upper bounds the the policy complexity, defined as the mutual information between states and actions. This is equivalent to a bound on the average codeword length. **(B)** The optimal solution prescribed by rate-distortion theory, featuring state-dependent *Q*(*s*, *a*) and state-independent *P*(*a*) components. The parameter β adjusts the relative contributions of the two components. **(C)** The reward-complexity frontier for an example task, derived by varying the β parameter in (B) and finding the corresponding optimal policies. (A-C) are adapted from (Lai and Gershman, 2024). **(D)** An example Huffman code with optimal codeword assignments. The code lengths are assigned based on the probability of decoding different symbols (leaf nodes). In policy compression, the codetree would be tailored to decode actions. **(E)** The same Huffman code visualized as a codetree. On the process level, we hypothesize that each bifurcation in the codetree manifests as an increase in neural representation dimensionality, which is required to separate the two possible branches downstream.

It is well known that context-dependent action selection incurs cognitive costs that affect human behavior (Zenon et al., 2019; Shenhav et al., 2017; Sayalı et al., 2023). For a resource-rational agent, we formalize the cognitive cost as the mutual information between states and actions, which we call the *policy complexity*:

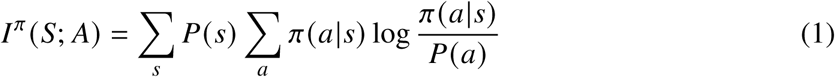

where *P*(*s*) is the state probability distribution and *P*(*a*) = _*s*_ *P*(*s*)π(*a*|*s*) is the marginal probability of choosing action *a* under the agent’s policy π(*a*|*s*). Intuitively, high-complexity policies preserve state information (e.g., deterministic mappings from states to actions) whereas low-complexity policies discard state information (e.g., random actions). One can additionally decompose policy complexity into individual *policy cost* terms for each state-action pair, 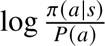, whose trial-wise average defines the policy complexity.

If the agent has infinite cognitive resources, it would be optimal to map each state to the most rewarding action. However, we assume that policies are subject to a capacity constraint *C* (a concept frequently invoked in the cognitive control literature; see (Botvinick and Cohen, 2014)), which, in our formulation, acts as an upper bound on policy complexity. The value of *C* can be voluntarily set by the agent (Liu et al., 2024), or, under cognitive load or response deadlines, limited by the cognitive resources available (Lai and Gershman, 2024; Liu and Gershman, 2025). In this formulation, for a given task with state probability distribution *P*(*s*) and state-specific action rewards *Q*(*s*, *a*), agents would maximize trial-averaged reward *V*^π^ = _*s*_ *P*(*s*) _*a*_ π(*a*|*s*) *Q*(*s*, *a*) subject to their constraint *I*^π^ (*S*; *A*) ≤ *C*. We can express this constrained optimization problem in Lagrange form:

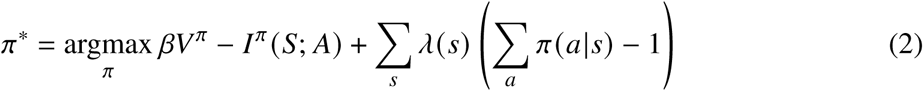

where β ≥ 0, λ(*s*) ≥ 0 ∀*s* ∈ *S* are Lagrange multipliers to ensure *I*^π^ (*S*; *A*) ≤ *C* and proper policy normalization: *I*^π^(*a*|*s*) = 1. Solving the Lagrange form yields the following optimal policy:

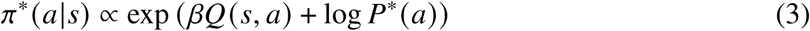

where *P*^∗^ (*a*) = _*s*_ *P*(*s*) π^∗^ (*a*|*s*) is the optimal marginal action distribution, and β is a Lagrange multiplier whose value depends on *C* in a task-specific manner. Despite the recursive nature of Equation 3, one can numerically find the optimal policy π^∗^ (*a*|*s*) by iterating updates of Equation 3 and the defining equation of *P*^∗^ (*a*) for different β values. This numerical process is known as the the Blahut-Arimoto algorithm, which is guaranteed to converge (Blahut, 1972; Arimoto, 1972; Tishby and Polani, 2010).

Intuitively, at high policy complexity (corresponding to large *C*), the value of β is large and the optimal policy is dominated by *Q*-values, which renders it state-dependent. At low policy complexity (small *C*), the value of β is close to 0 and *Q*-values have minimal impact on the optimal policy. Moreover, low-complexity policies are dominated by the log *P*^∗^ (*a*) term, which manifests as perseveration toward more frequently chosen actions in a state-independent manner (Figure 1B). In general, high-complexity policies enable more trial-averaged reward than low-complexity policies due to their state-dependence. By varying β and calculating the optimal policy, we can trace out the reward-complexity frontier, which delimits the maximal trial-averaged reward obtainable for a given policy complexity (Figure 1C).

### Connections to habits and perseveration

The formation of default actions or habits—and the cognitive cost required to override them—has been a central topic in cognitive control (Miller and Cohen, 2001; Shenhav et al., 2017; Zenon et al., 2019; Koechlin and Summerfield, 2007). The policy compression framework offers a resource-rational account of such habitual behavior: when cognitive resources are limited, biasing action selection toward frequently chosen actions is part of the optimal strategy. This arises because the optimal policy depends on the marginal action distribution *P*^∗^ (*a*). When *P*^∗^ (*a*) is uniform, the optimal policy simplifies to the standard softmax choice rule, π(*a*|*s*) ∝ exp[β*Q*(*s*, *a*)] (Sutton and Barto, 2018). However, when *P*^∗^ (*a*) is biased—reflecting that some actions have been chosen more frequently than others—the optimal policy will favor these actions, making them appear as default choices controlling for differences in *Q*(*s*, *a*). As cognitive resources become more limited (low policy complexity and small β), this bias is amplified: *P*^∗^ (*a*) increasingly dominates Equation 2, leading to stronger perseveration toward default actions. Such perseverative tendencies are not predicted by traditional softmax models with variable noise levels, where the policy converges to randomness as noise increases. In recent work, we have identified perseverative signatures in human decision making that are consistent with policy compression but not predicted by the softmax model (Lai and Gershman, 2024; Liu et al., 2024; Liu and Gershman, 2025), supporting the framework’s relevance. Unlike previous accounts that attribute habits to model-free learning (Krueger and Griffiths, 2018; Pauli et al., 2018), policy compression explains habits as value-independent tendencies shaped purely by past action frequencies. This perspective aligns with and provides a normative justification for recent findings suggesting that habits can form and exert influence independently of value representations (Zhang et al., 2024; Nebe et al., 2024; Miller et al., 2019).

### Connections to representational dimensionality

Recent studies of cognitive control have proposed a novel account of action selection, informed by the representational dimensionality of neural population codes. Representational dimensionality is defined as the minimum number of dimensions required to capture the variability of neural activity across task states (Ahlheim and Love, 2018; Fusi et al., 2016; Jazayeri and Ostojic, 2021; Badre et al., 2021). In this view, high dimensionality allows separating representations of different states, which enables state-dependent action selection. Conversely, low-dimensional representations reduce such separability, limiting the ability to tailor action selection to the current state (Badre et al., 2021; Badre, 2024). Specific to rule-based decision making, the benefit of high dimensionality manifests as the formation of conjunctive state representations integrating stimuli and rules. Consistent with this view, empirical work in cognitive control has observed transient increases in neural representational dimensionality during action selection, coinciding with higher-quality rule-based decisions (Kikumoto et al., 2022, 2024a).

The above representational account has rarely been linked to the information-theoretic view of cognitive control, which typically emphasizes top-down modulatory architectures instead (Koechlin and Summerfield, 2007; Fan, 2014; Badre et al., 2021). However, a theoretical bridge emerges when we consider algorithmic-level entropy codes inspired by information theory. To illustrate the connection, let us consider representing states as binary codewords generated through entropy coding, with the Huffman code (Huffman, 1952) as a canonical example (Figure 1D). In this scheme, each state codeword maps to a specific action by traversing a binary tree structure. State-dependent action selection requires that codewords corresponding to different states be sufficiently distinct to reach different leaf nodes. This necessitates longer codewords for readout, which in turn requires traversing more bifurcations in the codetree (Figure 1E). In the regime of errorless transmission, the policy’s complexity (in bits) corresponds to the average codelength, under an optimal entropy coding that minimizes this quantity (Cover, 1999). By adopting this entropy coding view of policy compression, we hypothesize that the codetree bifurcations during readout map onto transient increases in neural representational dimensionality. Consequently, behavioral policy complexity should reflect the representational dimensionality required to support the policy. Furthermore, under strict response deadlines that prevent full traversals of the codetree (as in the cognitive control dataset analyzed), the agent would be cut off at earlier bifurcations, reducing the state-dependence of their action selection and thus their policy complexity.

### Connections to response times

It is well established that exerting cognitive control leads to longer RTs (Matsumoto and Tanaka, 2004; Kool et al., 2017). Through the lens of optimal entropy codes, policy compression rationalizes longer RTs under high-complexity policies. Specifically, executing a high-complexity policy entails reading longer state codewords on average, which requires additional time to traverse the codetree (Lai and Gershman, 2021). This viewpoint closely mirrors neuroscience studies showing that transient increases in representational dimensionality are temporally extended during decision tasks, and that forceful cutoffs at an earlier timepoint diminishes rule-based performance (Kikumoto et al., 2024a). Similar information-theoretic explanations for RTs have long been applied to decision tasks that vary the number of available actions, encapsulated in the Hick-Hyman Law. This law formalizes the empirical observation that RT increases logarithmically with the number of possible actions, or equivalently, linearly with the amount of information transmitted (Hick, 1952; Hyman, 1953). Whether policy complexity similarly predicts RTs in cognitive control tasks remains an open empirical question.

## Materials and Methods

To directly test the predictions of policy compression for cognitive control tasks, we reanalyzed data from a previous electroencephalogram (EEG) study (Kikumoto et al., 2024a). We briefly summarize the study methods here, and refer readers to the original paper for more details.

## Participants

Forty-two participants (27 female, mean age 22 years) were recruited. The recruitment followed procedures approved by the Human Subjects Committee at the RIKEN, and all participants gave informed consent. The sex and gender of participants were determined based on self-report. They all had normal or corrected-to-normal vision and had no history of neurological or psychiatric disorders. No statistical method was used to predetermine the sample size. After preprocessing the EEG data, one participant was removed due to excessive amounts of artifacts (i.e., more than 25% of trials; see EEG recordings and preprocessing for details).

## Behavioral task

Participants completed a rule-based action selection task. On each trial, the participant simultaneously sees one of three rules (“horizontal”, “vertical”, “diagonal”) and one of four stimuli (situated in a 2-by-2 matrix), randomly sampled with equal probability for each trial, yielding a total of 12 rule-stimulus combinations (i.e., “states”). Participants must choose one of four actions, arranged in a 2-by-2 matrix on a number pad (Figure 2A). The rule determines the correct stimulus-action mappings. For example, if the current rule is “horizontal” and the stimulus is on the top-left, the correct action is to press the top-right action key on the number pad, as this position is horizontally adjacent to the stimulus (Figure 2B).

**Figure 2.**
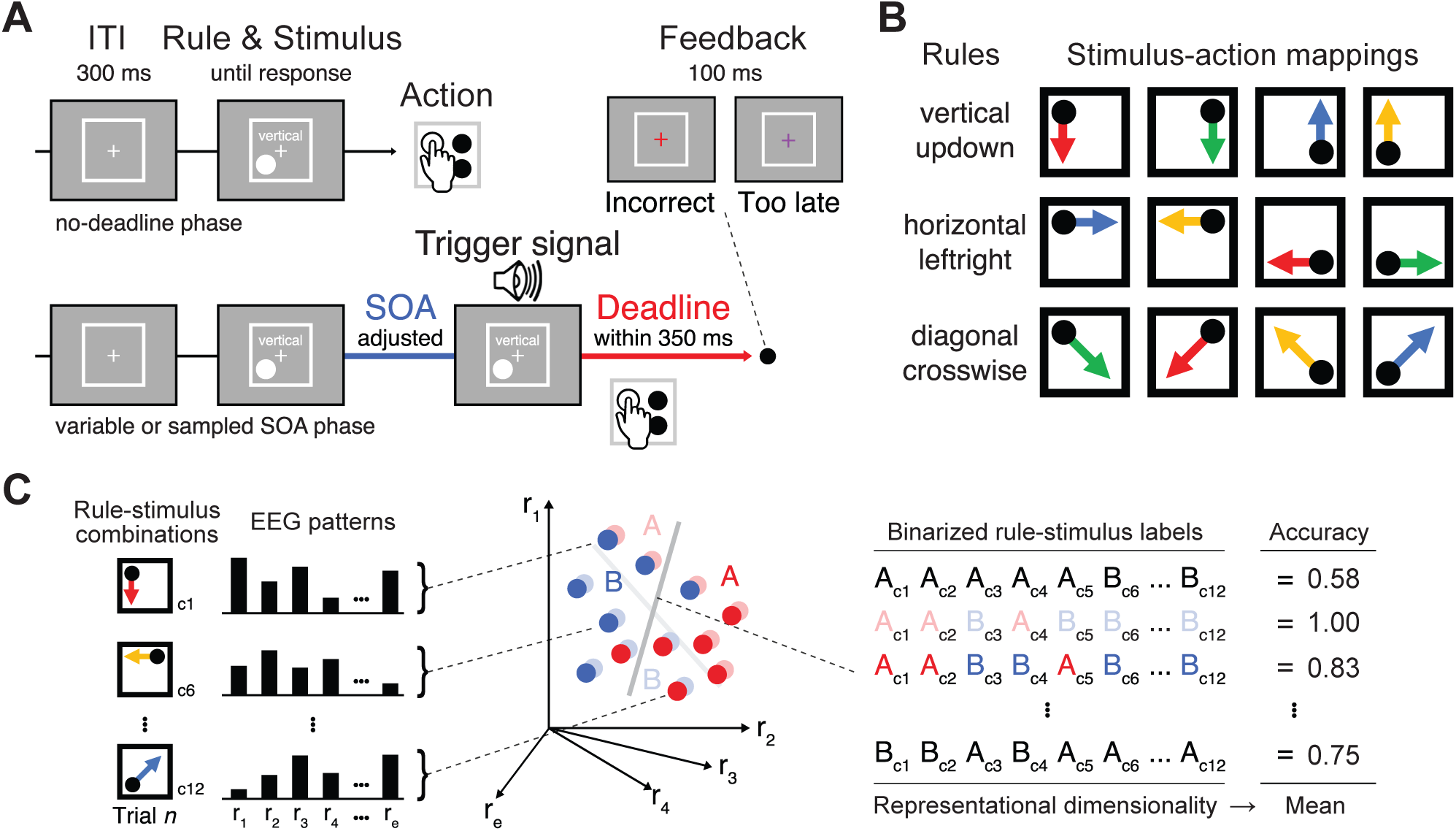
Behavioral task and EEG shattering dimensionality estimation. **(A)** Task pipeline and SOA conditions. On every trial, participants observe one randomly sampled rule and stimulus, and respond by choosing an action on the number pad. In the variable and sampled SOA phase, participants receive an auditory SOA signal some time interval after trial onset, and they must respond within 0-350ms after the signal’s presentation. In the variable SOA phase, the SOA intervals are determined for each subject separately to achieve three predefined accuracy levels. The SOA condition for each trial is sampled randomly, without block-level regularity that allows participants to adjust their strategy beforehand. **(B)** The stimulus-action mappings are determined by high-level rules. There are 3 possible rules, 4 possible stimuli (hence 3 × 4 = 12 rule-stimulus combinations, randomly sampled for each trial), and 4 possible actions. **(C)** Shattering dimensionality estimation. EEG signals (31 electrodes × 5 frequency bands = 155 dimensions) are fed into a linear decoder, to decode each of the (2^12^ − 2) = 4096 binary partitions of true rule-stimulus identity for each trial. The resulting trained decoder accuracy rates are used to estimate dimensionality of the underlying EEG representation. Adapted from Kikumoto et al. (2024a).

Trials were organized into experimental blocks, each lasting 18 seconds. For each block, participants were instructed to complete as many trials as possible. Trials that were initiated within the 18s block duration but extended beyond it were allowed to finish.

Blocks were further organized into task phases. Participants completed three task phases in order: the no-deadline phase, the variable stimulus-onset asynchrony (SOA) phase, and the sampled SOA phase. The no-deadline phase contained 25 blocks, where participant RTs are used to adjust the SOA interval for later phases.

In the variable SOA phase containing 35 blocks, for every trial, participants received an auditory signal some time after the trial onset. Upon this SOA onset, they were required to select an action within 350ms after hearing the auditory signal. If participants made a response before SOA onset or after 350ms after SOA onset, the trial response is considered invalid, and participants receive no reward. The time from trial onset to auditory signal presentation is called the SOA interval, which could take on 12 values determined for each participant. In each trial, the SOA interval was randomly sampled.

In the sampled SOA phase containing 185 blocks, SOAs were still present after trial onset, but each participant only received three unique SOA intervals, determined in a participant-specific fashion to induce accuracy rates of 50, 70 and 90% across trials (both invalid and incorrect responses are counted as inaccurate). The SOA interval was randomly sampled for every trial and thus interleaved. These three SOA intervals constitute task conditions and are labeled “short”, “medium”, and “long SOA” respectively. EEG data were collected in this phase.

### Estimation of SOA functions

To separately analyze the three accuracy levels during the sampled SOA phase, we preprocessed each participant’s RTs during the no-deadline and variable SOA phases. For the no-deadline phase, we computed each participant’s RT distribution, excluding trials with RTs slower than 5 SDs. We model the RT distribution using the ex-Gaussian distribution using the exgauss toolbox in Matlab 2019B.

In the variable SOA phase, each participant receives 12 possible SOA interval values. The values were determined based on deviations from the participant’s mean RT in the no-deadline phase (estimated as the mean of the fitted ex-Gaussion distribution). The deviations range from -450ms to +200ms with 50ms increments. Negative RTs were dropped during estimation. This allowed us to construct SOA functions that map SOA interval values to accuracy rates.

We modeled SOA functions using an exponential function: *p*(correct) = λ(1 − exp (−β(*t* − Ω)) if *t* > Ω, and *p*(correct) = 0 otherwise. The parameters (λ, β, Ω) were estimated by a nonlinear least square solver via the lsqcurvefit function in Matlab. The fitted SOA functions allowed us to identify SOA interval values that would produce particular accuracy levels (see (Kikumoto et al., 2024a) for visualizations of fitted SOA functions).

### EEG recordings and wavelet analysis

EEG recordings were collected during the variable SOA phase. The EEG signals were recorded using a Brain Products actiCHamp recording system (Brain Products GmbH), featuring 31 electrodes from a broad set of scale sites. The scalp EEG and EOG were amplified with an SA Instrumentation amplifier with a bandpass of 0.01-45Hz, and signals were downsampled at 250Hz using EEGLab93. We aligned trial-wise EEG recordings, spanning -800ms to +200ms relative to action onset (0ms).

After obtaining trial-wise EEG recordings, we decomposed them into 5 frequency bands via complex wavelet analysis, yielding a power measure for each timestep and frequency band. This produced 155 EEG features (31 electrodes × 5 frequency bands) for every participant, trial, and timestep (each lasting 1s/250Hz = 4ms). For more details on EEG recordings and wavelet analysis, please see Kikumoto et al. (2024a).

### Estimation of EEG representational dimensionality

To characterize neural representational dimensionality, we computed a shattering dimensionality estimate informed by the number of binary partitions of rule-stimulus combinations that are linearly separable based on the underlying neural representation (Rigotti et al., 2013; Bernardi et al., 2020; Courellis et al., 2024; Kikumoto et al., 2024a).

The EEG instantaneous power vectors (31 electrodes × 5 frequency bands) were further processed before decoder training. First, trials where responses occurred before SOA onset and trials where responses were completely omitted were excluded. Further, all trials in the first block (of the sampled SOA phase) were excluded. The remaining instantaneous power data was further averaged into 20ms time bins. For each participant and frequency band, the input vector entries were z-transformed across electrodes to remove effects that scaled all electrodes uniformly.

Based on how shattering dimensionality is typically estimated, we trained linear decoders on the EGG instantaneous power vectors to recover information on the trial’s presented rule-stimulus combination. Specifically, for each participant and 20ms time bin, we label each underlying trial’s presented rule-stimuli combination. We then bi-partition the 12 possible combinations, leading to (2^12^ − 2 = 4096) possible binary partitions (each of the 12 combinations may be included in or excluded from Group 0; minus the two binary partitions leading to all 0-labels or 1-labels). Thus, for each binary partition, we obtain a dataset over trials where the input consists of the EEG instantaneous power vector (5 frequency-bands × 31 electrodes = 155 dimensions), and the output are 0/1 trial-specific binary labels (Figure 2C left). For each of the 4096 binary partitions, we train a linear decoder to recover the binary labels from the input vector (Figure 2C center). This leads to a decoding accuracy value for the specific participant, time bin, and binary partition. The above process is repeated under repeated five-fold cross-validation, where the folds themselves were repeatedly partitioned 5 times through a random process. The resulting decoding accuracies are averaged.

After obtaining repeated, cross-validated decoding accuracies for each of the 4096 binary partitions, we further averaged over the decoding accuracies over binary partitions. This *partition-averaged decoding accuracy* measures the linear separability of neural representations across all possible binary partitions. Consistent with past work (Bernardi et al., 2020; Courellis et al., 2024), we use these partition-averaged decoding accuracies to construct a proxy of the EEG signals’ neural representational dimensionality (Figure 2C right). Specifically, the above partition-averaging process was done separately for each SOA task condition, leading to condition-specific representational dimensionalities for each participant and timepoint. We then aggregated the partition-averaged decoding accuracies over timepoints to compute the mean decoding accuracy across time bins for a trial, and additionally the trial-averaged mean of the maximum decoding accuracy over each trial’s time bins. The latter measure is our proxy for representational dimensionality.

Note that unlike some previous studies (Bernardi et al., 2020; Courellis et al., 2024), the rule-stimulus combinations used in this task did not feature strict dichotomies. Consequently, we decided to use a slightly different procedure in computing decoding accuracies, incorporating all (2^12^ − 2) possible binary partitions. While many of these binary partitions induce classification imbalance (e.g., all but one rule-stimulus combination being assigned the label 0), they do not pose a significant problem for our subsequent analysis. This is because the imbalance is present for all participants, SOA conditions, trials, and timesteps, and we only focus on comparing relative differences across partition-averaged decoding accuracies.

### Estimation of policy complexity

We defined policy complexity as the mutual information between the observed states and chosen actions. Following prior work (Gershman, 2020; Lai and Gershman, 2024; Liu et al., 2024), we estimated the policy complexity of each participant in each SOA condition using the Hutter estimator (Hutter, 2001). Specifically, for each of the 12 states (rule-stimulus combinations), we assume a symmetric Dirichlet prior with α = 0.1 for all actions chosen, and use the empirical action counts to reach a posterior Dirichlet distribution over action probabilities. We then estimate policy complexity as the mutual information of the posterior mean policy. The above procedure is informed by previous literature, reporting that the resulting estimates exhibit reasonably good performance when the joint distribution is sparse (Archer et al., 2014). The choice of α = 0.1 is informed by rate-distortion theory, stating that empirical trial-averaged reward values cannot be above the reward-complexity frontier. We have chosen α = 0.1 empirically to obey this property.

We also computed policy cost values 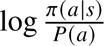 for individual trials. Given the blocked structure of the experiment, as well as the dynamically evolving nature of *P*(*a*), we decided to track π(*a*|*s*) and *P*(*a*) in an online manner for each block separately.

### Statistical analysis and modeling

We assessed the behavioral predictions of policy compression using two-sided paired t-tests across the two most extreme SOA conditions (low versus high). Regarding trial-averaged RT, we quantitatively connect it to policy compression through the fitting of linear mixed-effects models (LME), using each participant-condition’s policy complexity (both fixed and participant-specific random effects) to predict corresponding trial-averaged RTs. The resulting LME model was compared to null models with only intercepts via the Bayesian Information Criterion (BIC).

For the EEG data, we connected policy compression to the shattering representational dimensionality measures derived according to previous sections. We only analyze responsive trials with available EEG data, focusing on the decoding accuracies of timebins from -800ms to 0ms aligned to action onset (i.e., discarding post-action decoding accuracies from 0ms to +200ms). The dimensionality measure—trial-averaged mean of the maximum decoding accuracy over each trial’s timebins—was of particular interest, due to its connection with policy complexity—the average readout codelength of the policy’s optimal entropy coding. On the single-trial level, we studied the time-averaged decoding accuracy and policy costs. The mean is taken over timebins instead of the maximum, due to the noisiness of trial-wise EEG data. We again fitted LME models that used behavioral policy compression or policy cost measures (fixed and participant-specific random effects) to predict dimensionality or trial-wise mean decoding accuracy. The fitted LMEs were compared against null models with only intercepts or order effects (e.g., trial number) using BIC.

### Perseveration analysis

Given the policy compression framework’s rationalization of perseveration behavior, we computed probabilities of repeating the previous trial’s action, reapplying the previous trial’s rule to the current trial’s stimulus, and choosing actions based on the previous trial’s stimulus and the current trial’s rule. The perseveration analyses is done in a block-wise manner, so as to filter out the influence of resting time between blocks. Before computing the above probabilities, we excluded all omitted trials (in which participants did not respond), all trials preceded by an omitted trial, and the first trials of every block. Given the rule application analysis, we also excluded trials in which the participant had chosen an action incorrect under every possible rule (i.e., choosing the action that has the same number-key location as the stimulus). The application of these criteria excluded 9113 (8.33%) responsive trials across participants.

## Results

### Experimental predictions

The policy compression framework makes the following predictions: longer SOA conditions should be associated with 1) higher policy complexity; 2) higher trial-averaged RT; 3) higher accuracy. Furthermore, 4) the change in RT should be largely explained by changes in policy complexity, such that an LME model using participant-specific policy complexity levels to predict RT should identify positive fixed effects and win model comparison against a null LME model. Predictions 1, 2, and 4 arise due to the RT implications of policy complexity as discussed in the framework’s introduction.

Prediction 3 derives from the reward-complexity frontier associated with the task, coupled with the normative assumption that participants should achieve maximally attainable trial-averaged rewards at their chosen policy complexity levels. Policy compression also predicts that 5) the action entropy *H*(*A*|*S*)—capturing choice variability under the same state—would increase, as informed by the optimal policies at different β levels. Regarding perseveration patterns, the framework predicts that 6) the probability of repeating the previous trial’s action, should increase. Given the hierarchical rule-stimulus structure of the task, we further postulate 7) increases in the probability of applying the previous trial’s rule on the current trial’s stimulus. These predictions arise from the *P*^∗^ (*a*) term in Equation 3, and the fact that this marginal probability distribution must be dynamically updated across trials.

Regarding representational dimensionality, the policy compression framework makes the following predictions: on the single trial level, 8) an LME model using policy costs to predict single-trial mean decoding accuracy should perform better than a null model, which either features only fixed and random intercepts, or additionally non-policy-compression predictors including block-order effects, trial-order effects, and SOA condition. On the trial-aggregate level, we should see 9) higher trial-averaged maximum decoding accuracy over timesteps (proxy for representational dimensionality) for longer SOA conditions, and 10) an LME using participant-condition-specific policy complexity values should predict representational dimensionality better than a null model with only intercept effects. These predictions derive from the connection of optimal entropy codelengths to representational dimensionality necessary for policy execution, as discussed in the earlier policy compression section.

### Behavioral results

Participants completed an average of 2518 ± 24.28 trials each (mean ± SEM), resulting in a total of 103256 responsive trials for subsequent analysis.

Participant behavior was closely aligned with the task’s reward-complexity frontier (Figure 3A). This suggests that participants have achieved near maximal trial-averaged reward as allowed by their policy complexity level, supporting the relevance of the policy compression framework for cognitive control tasks. However, we also observed systematic deviations from the frontier at low levels of policy complexity. These deviations suggest an inefficient use of cognitive resources, caused by suboptimal behavioral patterns that we will elaborate on below.

**Figure 3.**
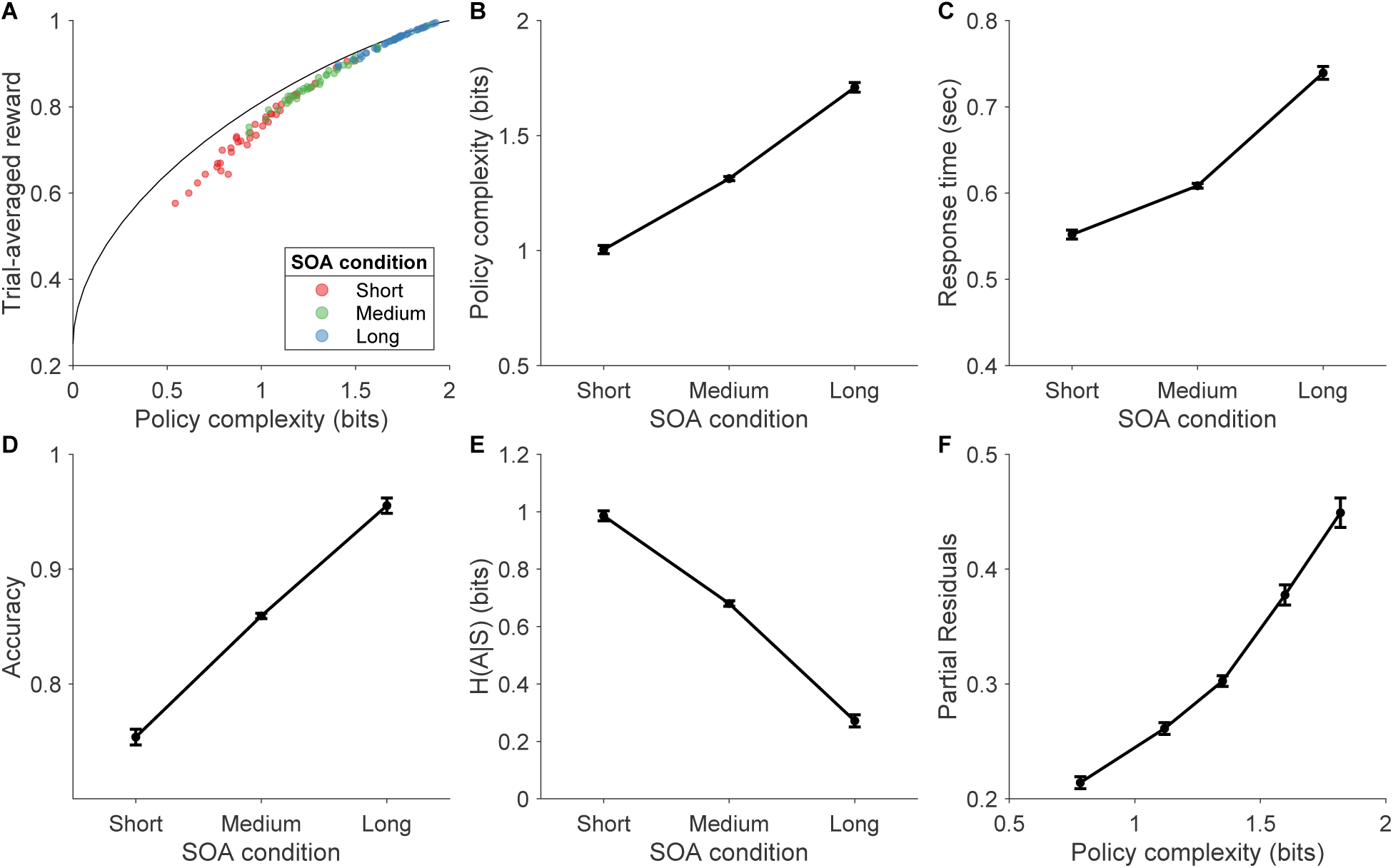
Behavioral results. Color denotes SOA condition. **(A)** Reward-complexity frontier and human data, where each of the 3 × 4 = 12 rule-stimulus combinations is treated as a separate state. Trial-averaged reward is binarized according to correct/incorrect responses provided, regardless of whether the response is valid (i.e. provided after the SOA onset but before another 350ms passed by). **(B)** Policy complexity, **(C)** RT, **(D)** Accuracy, **(E)** Action entropy *H*(*A*|*S*), **(F)** Partial residual plot for the fixed effects of the RT LME model, visualizing the effect of policy complexity on trial-averaged RT. Observations were binned into quantiles for visualization. For (A-E), error bars denote mean ± standard error of the mean (SEM) across participants (Cousineau, 2005); for (F), error bars denote mean ± SEM across binned observations.

The behavioral predictions of policy compression were predominately supported. In longer SOA conditions, participants adopted higher policy complexity (*t* (40) = −18.7, *p* < 10^−20^; Figure 3B) while incurring longer RTs (*t* (40) = −15.1, *p* < 10^−17^; Figure 3C), higher accuracy rates (*t* (40) = −15.2, *p* < 10^−17^; Figure 3D), and lower action entropy (*t* (40) = 19.1, *p* < 10^−20^; Figure 3E). To determine whether the increase in RTs was indeed associated with increased policy complexity, we fitted an LME model with fixed and random effects for both the intercept and policy complexity predictors to predict trial-averaged RTs. As expected, the fitted LME identified positive fixed effects for policy complexity (0.239 ± 0.0191, *t* (121) = 12.5, *p* < 10^−22^, random effects SD = 0.0891; Figure 3F), and outperforms a null LME model with only fixed and random effects for the intercept (ΔBIC = −121).

We next examined perseveration patterns as predicted by the framework. At the level of low-level actions, there was no evidence of increased perseveration: the probability of repeating the previous trial’s action did not significantly change across SOA conditions (*t* (40) = 0.289, *p* = 0.774; Figure 4A). In contrast, we observed prominent perseveration tendencies at the higher level. Under shorter SOAs, participants were more likely to reapply the rule they behaviorally followed on the previous trial to the current trial’s stimulus, suggesting rule-based perseveration (*t* (40) = 4.77, *p* < 10^−4^; Figure 4B). Additionally, they were also more likely to apply the current trial’s rule to the previous trial’s stimulus, which could be summarized as stimulus-based perseveration (*t* (40) = 2.75, *p* = 0.00889; Figure 4C).

**Figure 4.**
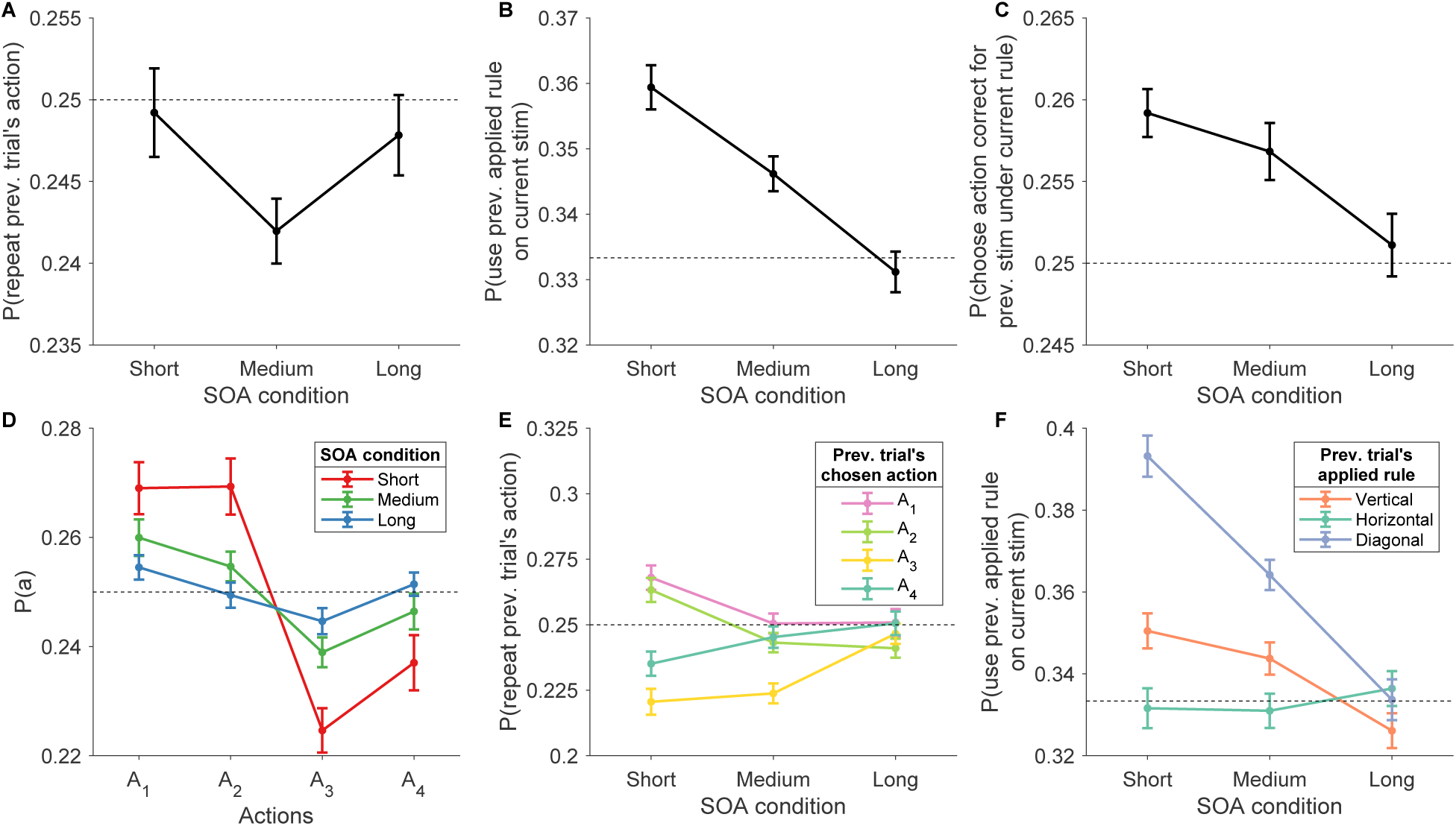
Perseveration. Color denotes different actions/rules. **(A)** Probability of repeating the previous trial’s action; **(B)** Probability of reapplying “the rule used by the agent on the previous trial” to the current trial’s stimulus. **(C)** Probability of choosing the previous trial’s stimulus’s correct action under the current trial’s rule. **(D)** *P*(*a*) across SOA conditions. **(E)** (A) stratified by action identity. **(F)** (B) stratified by rule identity. The dotted lines in each panel denote chance levels of action selection or rule application. Error bars denote mean ± standard error of the mean (SEM) across participants.

To better understand the empirical suboptimalities observed in Figure 3A and the lack of significant differences in Figure 4A, we conducted follow-up analyses stratified by action identities. The observed suboptimality at low policy complexity likely resulted from a biased marginal action distributions *P*(*a*). Specifically, in the short SOA condition, participants disproportionately favored two specific actions—*a*_1_ and *a*_2_, corresponding to the top two keys on the number pad (mean *P*(*a*) for *a*_1_ and *a*_2_ versus that for *a*_3_ and *a*_4_: *t* (40) = 5.78, *p* < 10^−6^; Figure 4D). This preference deviates from the framework’s normative predictions, which prescribe optimal policies with equiprobable marginal distributions *P*(*a*) across β values.

We conducted similar follow-up analyses of perseveration patterns, stratifying by either action or rule identity. We again observed the empirical preference for *a*_1_ and *a*_2_ in the short SOA condition (Figure 4E). Additionally, participants increasingly favored the diagonal rule under shorter SOAs (Figure 4F).

### EEG dimensionality results

On the single-trial level, we assessed whether policy cost—whose average across trials defines the policy complexity—predicts trial-wise mean EEG decoding accuracy. The corresponding LME yielded positive fixed effects for policy cost (0.000887 ± 0.000177, *t* (96630) = 5.01, *p* < 10^−6^, random effects SD = 0.000500; Figure 5A), while also outperforming a null model containing only fixed and random effects on the intercept (ΔBIC = −16.5874). To rule out potential confounds, we constructed alternative null models that included block order, trial order, and SOA condition as predictors, either individually or in combination. Model comparison consistently preferred the policy cost LME (ΔBIC ∈ [−138.1, −42.03] across all pairwise comparisons).

**Figure 5.**
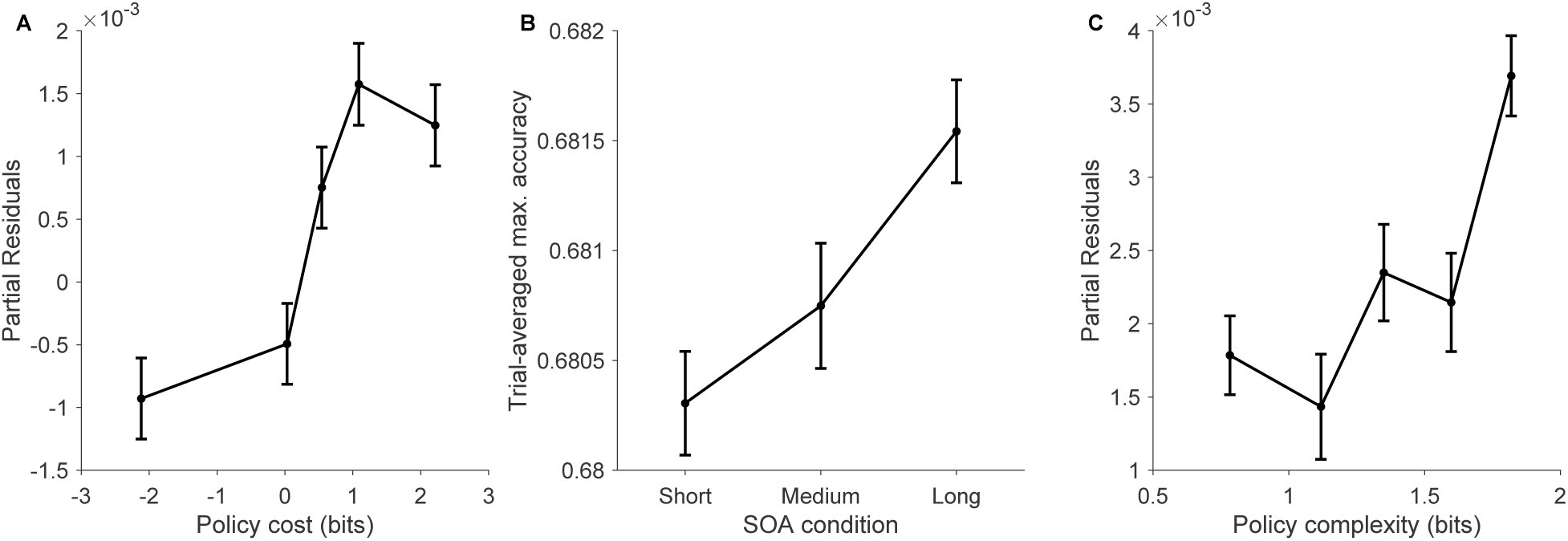
Neural dimensionality. **(A)** Partial residual plot for the fixed effects of the trial-by-trial LME, visualizing the effect of policy cost on decoding accuracy. Error bars reflect mean ± SEM across LME observations. **(B)** Mean ± standard error of the mean (SEM) of participant maximum dimensionality across SOA conditions. For each participant, the maximum decoding accuracy for each trial is computed over time, and then averaged across trials to reach the dimensionality measure. **(C)** Partial residual plot for the fixed effects of the trial-aggregated LME, visualizing the effect of policy complexity on the maximum dimensionality as computed in (B). Error bars denote mean ± SEM across LME observations. For (A) and (C), LME observations were binned into quantiles for visualization.

At the trial-aggregated level, we examined whether policy complexity predicts representational dimensionality, measured by the trial-averaged maximum decoding accuracy across timesteps. As predicted, representational dimensionality was higher for longer SOA conditions (*t* (40) = −3.31, *p* = 0.00198; Figure 5B). Additionally, an LME model predicting representational dimensionality from policy complexity revealed positive fixed effects for the latter (0.00171 ± 0.000576, *t* (121) = 2.96, *p* < 0.00365, random effects SD = 0.000335; Figure 5C). However, in contrast to the single-trial results, the above model lost model comparison against a null model with only fixed and random effects for the intercept (ΔBIC = 5.73), indicating ambiguous evidence for the relationship at the trial-aggregate level.

## Discussion

In this paper, we introduced policy compression as a normative framework for understanding cognitive control. Extending its previous applications to multiarmed bandit tasks, we conceptualized the core challenge of cognitive control—dynamically selecting actions based on context-specific goals (Badre, 2024; Fan, 2014)—as a constrained optimization problem: agents must maximize external rewards under cognitive resource constraints. We formalized these resource constraints using policy complexity, defined as the mutual information between environmental states and the policy-assigned actions. This formulation captures the informational cost of context-sensitive decision making, as postulated by previous accounts (Fan, 2014; Zenon et al., 2019; Shenhav et al., 2017).

We demonstrated the theoretical utility of policy compression by linking it to several phenomena central to cognitive control. This includes the emergence of default actions and habits, the effortful overriding of habitual responses, the neural representational dimensionality required to support control, and the prolonged response times associated with increased context-sensitivity. In doing so, policy compression offers a unifying explanation for these diverse observations. Notably, our approach advances traditional information-theoretic views of cognitive control (Koechlin and Summerfield, 2007; Fan, 2014) by adopting a resource-rational perspective that explicitly incorporates cognitive costs (Shenhav et al., 2017), and by grounding the theory in recent neuroscientific findings on representational geometry and dimensionality (Bernardi et al., 2020; Rigotti et al., 2013; Kikumoto et al., 2024a).

To evaluate the behavioral and neural predictions of policy compression for cognitive control tasks, we analyzed a previously collected dataset featuring rule-dependent action selection and EEG recordings. Our behavioral results support the framework’s core predictions: across SOA conditions, behavior was close to the optimal reward-complexity frontier, exhibiting varying levels of policy complexity that explain corresponding changes in response times and action stochasticity. In contrast, the framework’s predictions on perseveration are partially supported. While action-level perseveration did not significantly change across conditions (which could result from a suboptimal over-reliance on two of the four actions), participants demonstrated a marked tendency to repeat previously applied rules or stimuli, suggesting compression at higher levels of abstraction.

Combined with previous studies on multiarmed bandits, the above behavioral findings contribute to a growing body of work suggesting that policy compression provides a domain-general account of context-sensitive behavior. A limitation of this prior work is the requirement that participants learn their policy from feedback; the policy compression framework is fundamentally about limitations on action selection, not learning, and thus learning dynamics obscure the framework’s predictions. Cognitive control tasks—where learning is deliberately minimized by the task design—offer a cleaner test of the theoretical predictions. Our results suggest that human behavior, in the absence of reward learning, aligns even more closely with the framework’s optimality predictions. These promising findings encourage future work connecting policy compression to neighboring topics in cognitive control, such as task-switching costs (Zenon et al., 2019) and aversion to hierarchical task abstraction (Sayalı et al., 2023), both of which can be interpreted as consequences of limited information-processing capacity.

More behavioral work is needed to further assess policy compression’s predictions for cognitive control tasks. First, the *P*^∗^ (*a*) term in Equation 3 could be operationalized as repeating previous actions as we hypothesized, but it is fundamentally a bias towards more frequently taken actions. The current dataset did not feature manipulations over the reward structure *Q*(*s*, *a*), which, when present, could induce changes in the optimal marginal action distribution *P*^∗^ (*a*) by biasing it towards certain low-level actions or high-level rules (Lai and Gershman, 2024; Liu et al., 2024; Liu and Gershman, 2025). Novel task designs that incorporate such manipulations would allow us to test whether and how human adaptively compress their policies in response to environmental regularities.

Second, our current policy formulation is limited in that it groups both rules and stimuli into a joint state representation. However, participants demonstrate different perseveration patterns at action and rule levels, featuring significant inter-conditional differences in one but not the other. One possible explanation is that different levels of the control hierarchy are subject to distinct information-processing bottlenecks (Sayalı et al., 2023)—an idea that merits further investigation. To address this limitation, we plan to develop a hierarchical extension of the policy compression that models rule application and action selection as a two-stage process with separate information bottlenecks.

Third, the usage of SOA conditions may delay matured responses, as participants are discouraged from responding before SOA onset. However, it is unlikely that the current RT results stem solely from this delaying effect, as we have observed significant changes in policy complexity, accuracy, and action entropy across SOA conditions. The above phenomenon could not be explained by the delaying account, which would predict similar behavioral statistics despite differences in RT. To further mitigate delay effects, we have repeated the RT regression analysis for the short SOA condition only, which is least prone to relevant issues. As predicted by the framework, the regression model featuring policy complexity again yielded positive effects (0.150±0.0340, *t* (39) = 4.18, *p* < 0.001), and won against a null model containing only the intercept (ΔBIC= −10.8). Future empirical studies could remove this confound by replacing SOA deadlines with RT deadlines common to policy compression studies (Lai and Gershman, 2024; Liu and Gershman, 2025), which would similarly compel responses but never delay them.

While previous studies have already identified neural correlates of policy cost in striatal dopamine (Gershman and Lak, 2025) and hypothalamic hypocretin/orexin (Tesmer et al., 2025), the initial site of policy cost computation, and how it is used by downstream areas, remains unclear. In addition, previous studies have focused on assessing the existence of policy cost representations in the brain, without providing a process-level account of how such costs were derived in the first place.

Our contribution here is introducing neural representational dimensionality as a cortical correlate of policy compression, thereby connecting it to the information-theoretic view of cognitive control. We found that greater decoding accuracy over binary state label partitions—a trial-level proxy for increased neural dimensionality—was correlated with higher policy cost. This finding aligns with the entropy coding perspective, where high-dimensional, fine-grained representations are required for supporting highly state-dependent policies. Beyond single-trial correlations, we also observed significant differences in neural representational dimensionality across SOA conditions. Although the relationship was relatively weak, the observed trends encourage future experiments specifically designed to test the neural-level predictions of policy compression.

The neural correlates of policy complexity is thus a large topic that warrants additional investigation. One prominent future direction entails developing biologically plausible neural circuits for implementing near-optimal entropy codes. Combined with causal intervention studies, such models could further inform how the brain may implement policy compression. Another potential direction involves examining the development and plasticity of these high-dimensional neural representations. While policy compression itself does not prescribe a specific learning algorithm, studying how neural representations gradually attain optimality through practice (Kikumoto et al., 2024b) may help explain human deviations from the normative predictions of policy compression.

## Author contributions statement

Project conception: S.L. and S.G.; Experiment conception: A.K.; Experiment methodology: A.K.; Data Collection: A.K.; Analysis methodology: S.L. and S.G.; Analysis and Interpretation: S.L.; Manuscript original draft: S.L.; Manuscript edits: A.K., D.B., and S.G. Experiment funding acquisition: D.B.; Project supervision: S.G..

